# Influence of Physicochemical Parameters on the in vitro Stability of DNA Tetrahedral Nanostructures

**DOI:** 10.64898/2026.05.10.724064

**Authors:** Janvi Viroja, Kiransinh Rajput, Sweny Jain, Dhiraj Bhatia

## Abstract

Tetrahedral DNA nanostructures (TDNs) are promising nanocarriers due to their structural precision, biocompatibility, and efficient cellular uptake. However, their stability under physiological conditions remains a key challenge. In this study, TDNs were synthesized via a one-pot thermal annealing method and characterized using native PAGE, dynamic light scattering (DLS), and zeta potential analysis, confirming uniform size (∼13 nm) and negative surface charge. Their stability was systematically evaluated across different biological media (DMEM complete, serum-free DMEM, and E3), temperatures (4 °C, 25 °C, and 37 °C), and pH conditions (4.0, 7.0, and 8.5) over 24 h. Results revealed rapid degradation in serum-containing medium, increased instability at higher temperatures, and reduced stability under acidic conditions, while serum-free, lower-temperature, and neutral to mildly basic environments enhanced structural integrity. These findings highlight the strong environmental dependence of TDN stability and provide insights for optimizing their design for biomedical applications.

## 1. Introduction

DNA is a biopolymer that stores and transfers genetic information and has increasingly emerged as a programmable material for nanotechnology applications due to its unique structural and chemical properties [1] [2]. The development of DNA nanotechnology, first introduced by Nadrian Seeman in the 1980s, enabled the rational design of stable nanostructures through sequence-specific Watson–Crick base pairing [3]. This advancement led to the construction of diverse two-dimensional and three-dimensional DNA architectures for applications in biosensing, bioimaging, drug delivery, and nanomedicine [5] [9]. Among the various DNA nanostructures, tetrahedral DNA nanostructures (TDNs) have gained considerable attention because of their simple design, structural rigidity, nanoscale dimensions, and excellent biocompatibility [14]. TDNs are typically assembled from four complementary single-stranded DNA oligonucleotides that self-assemble into a three-dimensional tetrahedral geometry. Owing to their compact structure, programmability, ease of functionalization, and efficient cellular uptake, TDNs have emerged as promising platforms for therapeutic delivery and biomedical applications [17] [18]. The structural stability of DNA nanostructures is highly dependent on environmental conditions, including nuclease activity, ionic strength, pH, temperature, and biological media composition [13]. Serum-containing environments have been reported to accelerate degradation because of enhanced enzymatic activity, whereas ion-balanced buffer systems improve structural preservation [16]. In addition, elevated temperatures and acidic pH conditions may destabilize DNA duplex interactions and promote nanostructure degradation [15]. Despite increasing interest in TDN-based biomedical applications, comprehensive evaluation of their stability under multiple biologically relevant conditions remains limited [11].

Therefore, in the present study, TDN stability was systematically evaluated under different biological media (DMEM complete medium, serum-free DMEM, and E3 medium), temperature settings (4 °C, 25 °C, and 37 °C), and pH environments (4.0, 7.0, and 8.5) over a 24 h period. Structural integrity and degradation behavior were analyzed using native polyacrylamide gel electrophoresis (PAGE) with normalized band intensity measurements. In addition, physicochemical characterization of TDNs was performed using dynamic light scattering (DLS) and zeta potential analysis to evaluate hydrodynamic size, surface charge, and colloidal stability. This study provides a comparative stability profile of TDNs under biologically relevant environments and provides valuable insight into the rational design and optimization of DNA nanostructures for future biomedical and nanotherapeutic applications.

## 2. Materials & Method

Specific DNA molecules (referred to as M1-M4) were bought from Sigma in the U.S. and purified with high-performance liquid chromatography (HPLC). They were dissolved in pure water from Thermo Fisher Scientific and stored at -20°C. Magnesium chloride (MgCl_2_) was obtained from Sigma-Aldrich, a part of Merck in the U.S. 25bp DNA step Ladder from PROMEGA Madison, WI USA.Dulbecco’s Modified Eagle Medium (DMEM) and serum-free DMEM were purchased from Gibco, a division of Thermo Fisher Scientific in the United States. The E3 medium was formulated using sodium chloride (NaCl), potassium chloride (KCl), calcium chloride (CaCl_2_), and magnesium sulfate (MgSO_4_) from Sigma-Aldrich in the United States. Phosphate-buffered saline (PBS) at pH 7.4 and Tris-acetate-EDTA (TAE) buffer were obtained from HIMEDIA in India. Acetate buffer at pH 4.0 and Tris-HCl buffer at pH 8.5 were formulated utilizing standard chemicals sourced from Sigma-Aldrich (USA) for pH-dependent investigations. Polyacrylamide gel electrophoresis supplies such as acrylamide/bis-acrylamide solution, ammonium persulfate (APS), and TEMED were acquired from Bio-Rad (USA). HIMEDIA (India) supplied agarose, 6X Gel DNA loading dye from GeNei (Bengaluru, India) and associated materials, while Bio-Rad provided ethidium bromide (EtBr).

### Instruments used

We used an electrophoresis system to do gel electrophoresis and looked at the gels under a ChemiDoc MP Imaging System by Bio-Rad. Malvern analytical Zetasizer Nano ZS instrument. To control the temperature, we used an Thermo Scientific incubator at 25°C, and 37°C. All liquids were handled with calibrated micropipettes.

## 2.1 Method

### 2.2 One-pot synthesis of DNA Tetrahedron

Singal stranded oligonucleotides M1, M2, M3, and M4 were prepared by dissolving them in nuclease-free water and heating them at 70°C for 1.5 hours at 350 rpm from their 100 μM stocks. For the experiment, a 10 μM solution was made by diluting these oligonucleotides with either 1X nuclease-free water. To produce DNA TD, equal amounts of M1, M2, M3, and M4 were used. To improve the stability of the DNA structure, 2mM of MgCl_2_ was added. This helps to reduce the repulsion between the DNA strands and make the structure more rigid. The process involved heating the oligonucleotides in a thermocycler to 95°C for 30 minutes, then gradually cooling it down in steps of 5°C to reach 4°C, with each step lasting 15 minutes. The resulting DNA tetrahedron (TD) was obtained at a concentration of 2.5 μM and stored at 4°C for later use.

### 2.3 Electrophoretic mobility shift assay (EMSA)

Using a 10% native polyacrylamide gel, the electrophoretic mobility shift assay (EMSA) was used to verify that the higher-order tetrahedral DNA nanostructure (TDN) had formed. The reaction mixture was made up of 1.5 μL of 6X loading dye and 10 μL of TDN sample. For 100 minutes, electrophoresis was performed at 80 V under native conditions. After electrophoresis stained with ethidium bromide for 10 minutes. The gel was seen using a Bio-Rad ChemiDoc MP Imaging System.

### 2.4 Dynamic Light Scattering (DLS) & zeta potential

To measure the hydrodynamic diameter of the formed nanostructure, DLS was performed. An aliquot of 2.5 μM TDN was diluted 1:10 with nuclease-free water and centrifuged at 10,000 rpm for 10 minutes. After collecting 950 μL of supernatant, it was adjusted to a final volume of 1 mL using nuclease-free water. The produced sample was then subjected to DLS and zeta potential analysis on a Malvern Zetasizer Nano ZS device. DLS was used to quantify the hydrodynamic diameter, and zeta potential measurements were taken to assess the surface charge and colloidal stability of the nanostructures. To ensure accuracy and reproducibility, all measurements were taken three times.

## 3. Results

### 3.1 TD synthesis & characterization

In a one-pot synthesis approach, use the thermal annealing protocol to create a DNA tetrahedron with three additional oligonucleotides (M2, M3, and M4). Confirmation was performed using EMSA. EMSA produced a ladder-like pattern as the number of oligonucleotides in the reaction increased. The bands’ mobility was similarly reduced, indicating the creation of a higher-order DNA nanostructure as the number of primers increased. (Figure 2 a). The average hydrodynamic size and the zeta potential of the formed nanostructure was measured using DLS. The hydrodynamic size of TD and is around 12.51± 3.2 nm (Figure 2 b). Zeta potential measurements showed that the native DNA tetrahedron (TD) possessed a surface charge of −14.11 mV, consistent with its negatively charged phosphate backbone. Upon conjugation with AT, the zeta potential shifted to +4.5 mV, confirming successful surface modification. This charge reversal suggests reduced electrostatic repulsion with the cell membrane and indicates enhanced potential for cellular uptake while maintaining acceptable colloidal stability (Figure 2 c).

**Figure 1:**
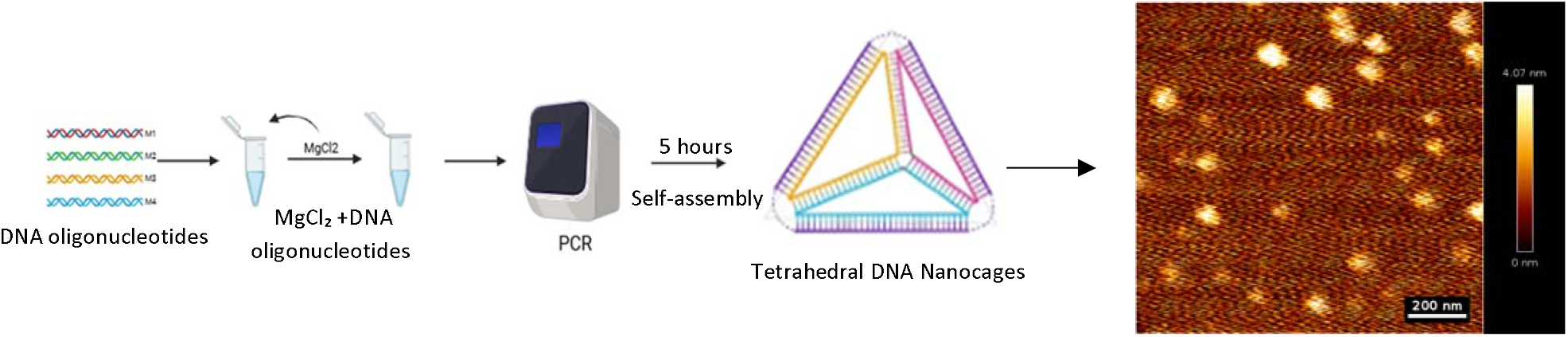
Schematic illustration of the one-pot synthesis and self-assembly of DNA tetrahedron nanostructures. Four single-stranded DNA oligonucleotides (M1–M4) were mixed in equal concentrations with 2 mM MgCl_2_, which reduces electrostatic repulsion between negatively charged DNA strands and enhances structural stabilization. The mixture was subjected to thermal annealing in a thermocycler, followed by gradual cooling to promote controlled self-assembly over 5 h, resulting in the formation of tetrahedral DNA nanocages. Representative image of AFM for DNA tetrahedron.

**Figure 2:**
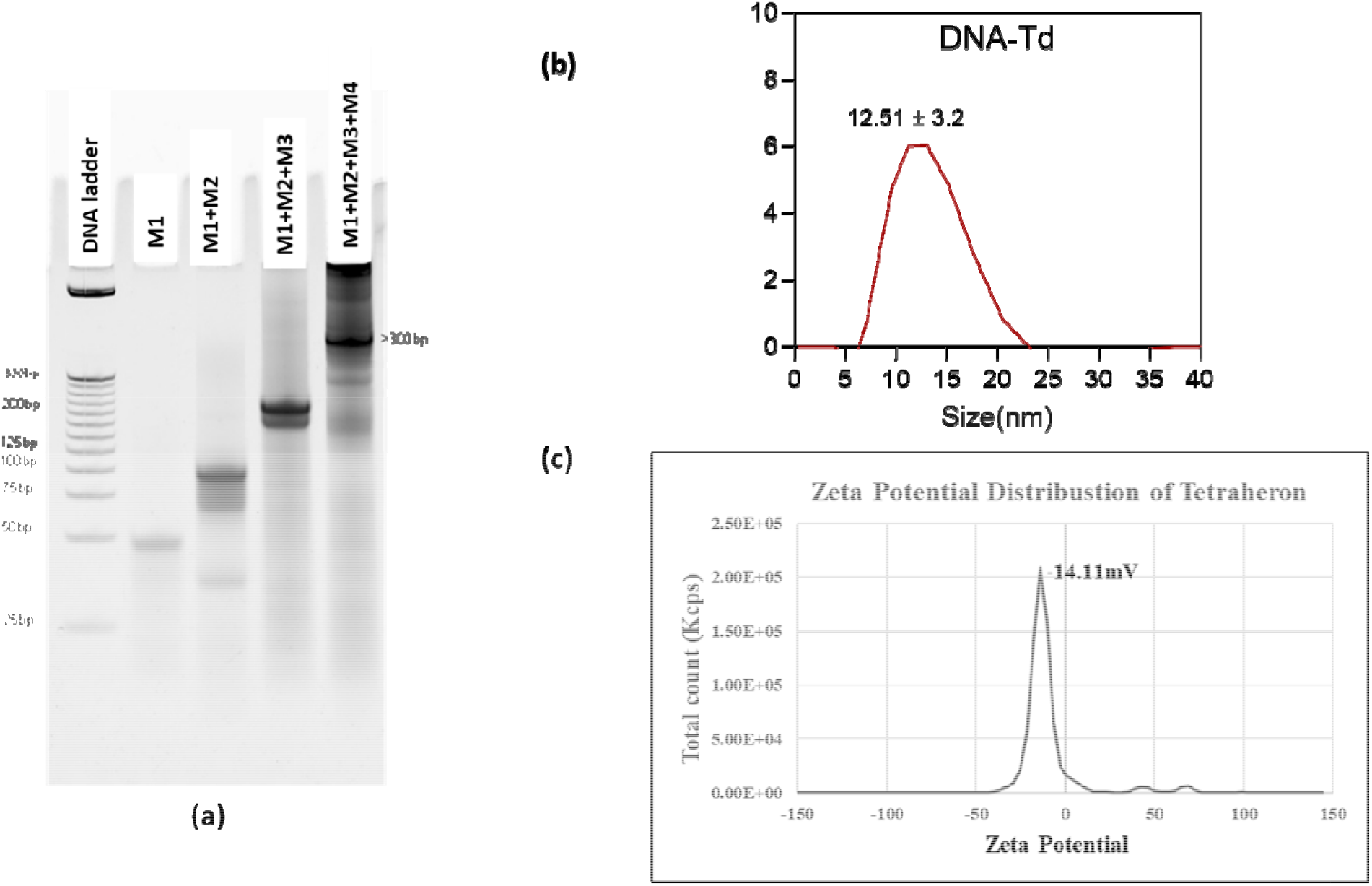
Characterization of DNA tetrahedron assembly and physicochemical properties. (a) Native PAGE analysis showing stepwise assembly of the DNA tetrahedron using individual strands (M1), partial assemblies (M1+M2 and M1+M2+M3), and the fully assembled structure (M1+M2+M3+M4). Progressive retardation in band mobility confirms successful hierarchical formation of the tetrahedral nanostructure. DNA ladder was used as the molecular size marker. (b) Dynamic light scattering (DLS) analysis showing the hydrodynamic size distribution of the DNA tetrahedron with an average particle size of approximately 12.51± 3.2 nm. (c) Zeta potential distribution of the DNA tetrahedron demonstrating a surface charge of approximately −14.1 mV, indicating colloidal stability of the assembled nanostructure.

### 3.2 Stability of DNA Tetrahedron under Different Environmental Conditions

The stability of DNA tetrahedron nanostructures was evaluated under different environmental conditions, including biological media, temperature, and pH, over 24 h using normalized band intensity (%). All samples showed comparable to control (∼100%).In different biological media (Figure 3), a time-dependent decrease in stability was observed. The structures in DMEM complete medium showed the fastest reduction in band intensity, indicating significant degradation, likely due to serum components. In contrast, serum-free medium exhibited improved stability with slower degradation, while E3 medium showed intermediate to high stability.

**Figure 3:**
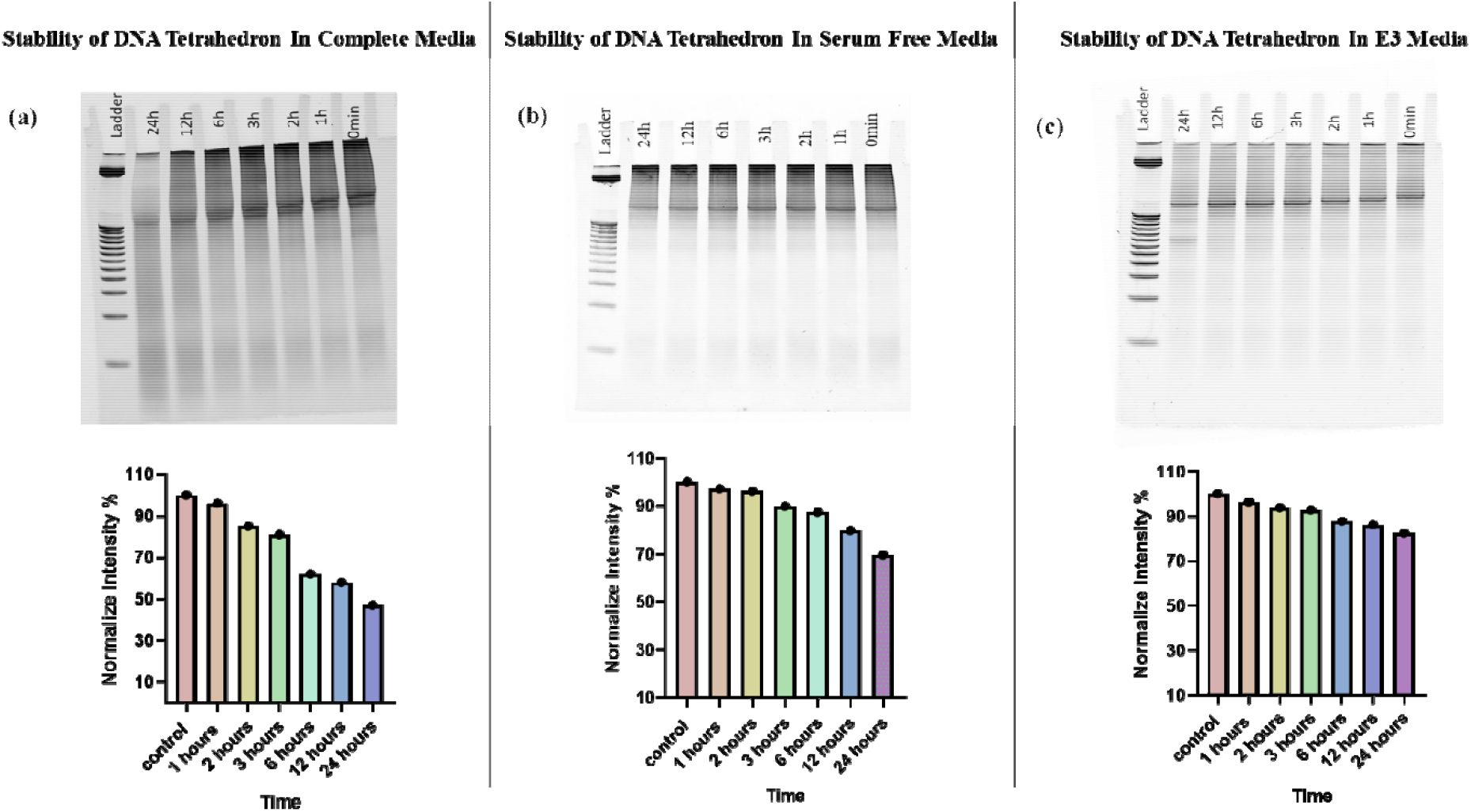
Stability analysis of DNA tetrahedron in different biological media over 24 h. Native PAGE images (top panel) and corresponding normalized band intensity profiles (bottom panel) of samples incubated in (a) DMEM complete medium, (b) serum-free medium, and (c) E3 medium.

Temperature-dependent analysis (Figure 4) revealed that stability decreased with increasing temperature. The structures remained highly stable at 4 °C, showing only a slight decrease in band intensity. At 25 °C, moderate stability was observed, whereas at 37 °C, a more pronounced reduction indicated increased degradation at higher temperature.

**Figure 4:**
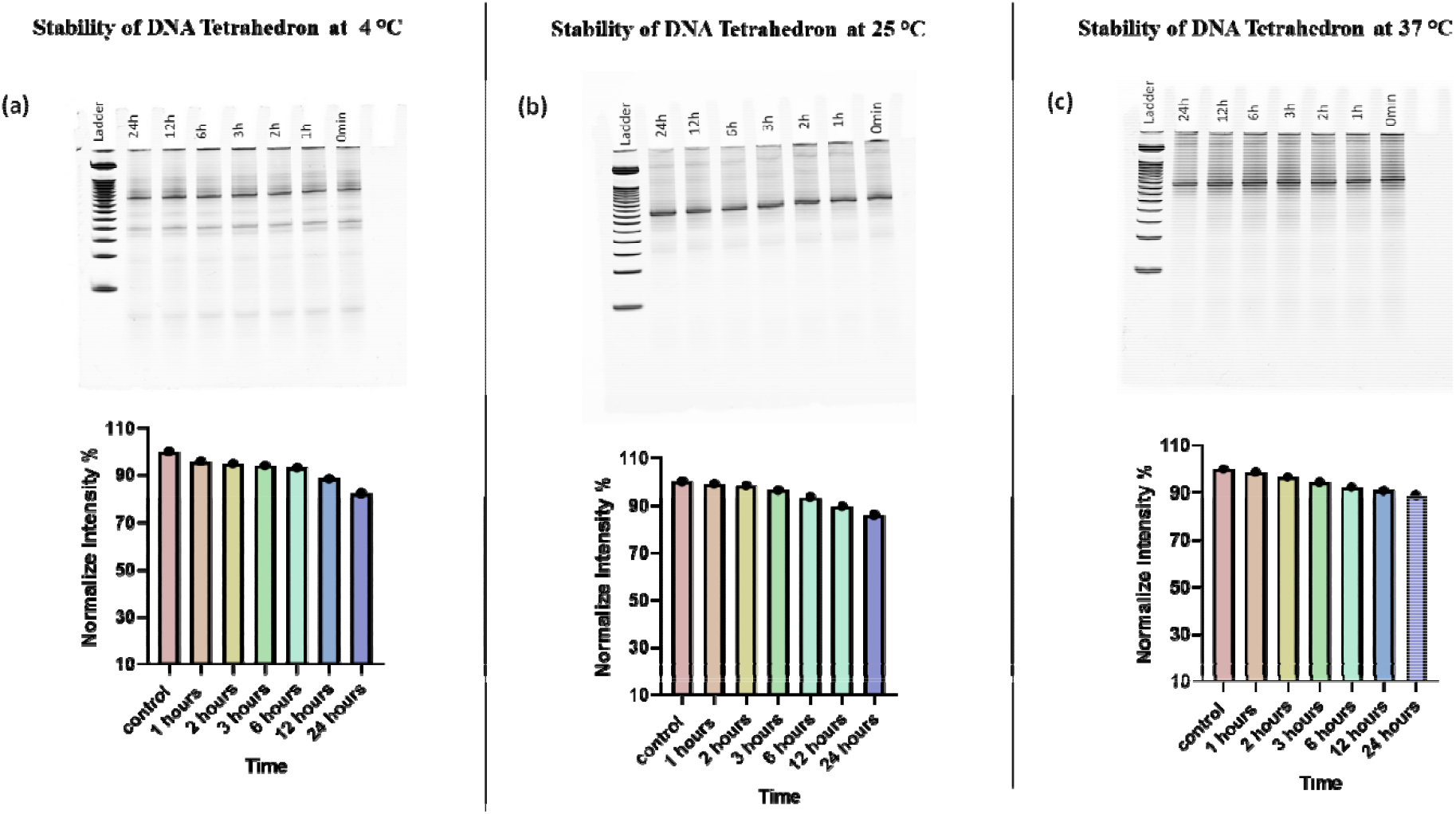
Temperature-dependent stability of DNA tetrahedron over 24 h. Native PAGE images (top panel) and corresponding normalized band intensity profiles (bottom panel) of samples incubated at (a) 4 °C, (b) 25 °C, and (c) 37 °C.

**Figure 5:**
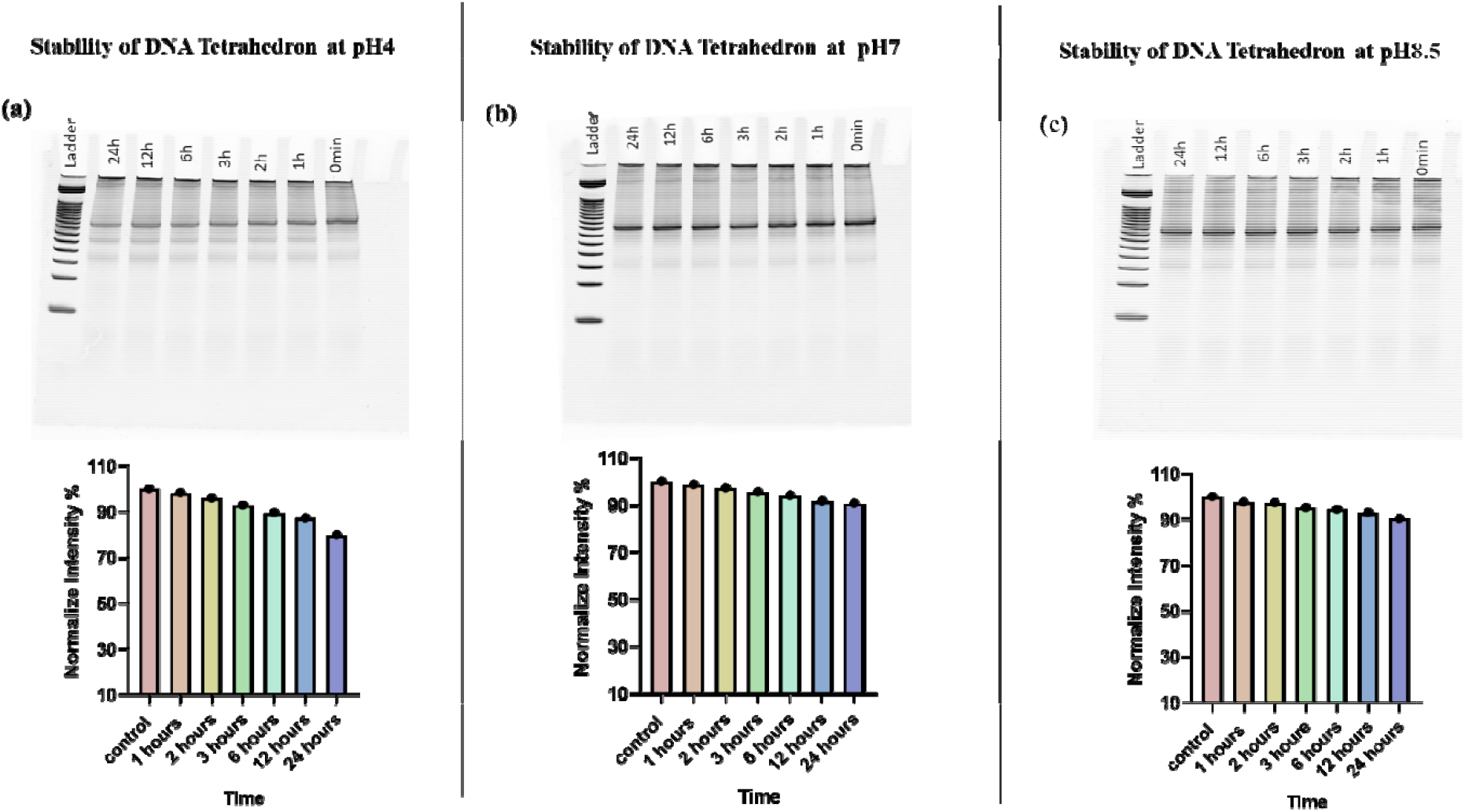
pH-dependent stability of DNA tetrahedron over 24 h. Native PAGE images (top panel) and corresponding normalized band intensity profiles (bottom panel) of samples incubated at (a) pH 4, (b) pH 7, and (c) pH 8.5.

Similarly, pH-dependent stability (Figure 2c) showed that acidic conditions (pH 4) led to faster degradation, as evidenced by a clear time-dependent decrease in band intensity. In contrast, neutral pH (pH 7) maintained high structural stability, while mildly basic conditions (pH8.5) showed moderate to good stability. Overall, these results demonstrate that DNA tetrahedron stability is strongly influenced by environmental conditions. The presence of serum, elevated temperature, and acidic pH accelerate degradation, whereas low temperature and neutral to slightly basic pH enhance structural stability.

### 3.2 Comparative Stability of DNA Tetrahedron under Different Conditions

The stability of DNA tetrahedron nanostructures was evaluated under different biological media, temperature, and pH conditions over 24 h using normalized band intensity (%). All samples showed comparable initial stability (∼100%). In different media (Figure 6a), a time-dependent decrease in stability was observed, with the fastest degradation in DMEM complete medium, followed by serum-free medium, while E3 medium maintained the highest stability. Temperature-dependent analysis (Figure 6b) showed that stability decreased with increasing temperature, with the fastest reduction at 37 °C, moderate stability at 25 °C, and highest stability at 4 °Similarly, pH-dependent stability (Figure 6c) indicated that acidic conditions (pH 4) resulted in the fastest degradation, while neutral (pH 7) and slightly basic conditions (pH 8.5) provided improved stability, with pH 8.5 showing the highest structural integrity. Overall, these results demonstrate that DNA tetrahedron stability is strongly influenced by environmental conditions, with serum components, higher temperature, and acidic pH promoting degradation, while controlled conditions enhance structural stability.

**Figure 6:**
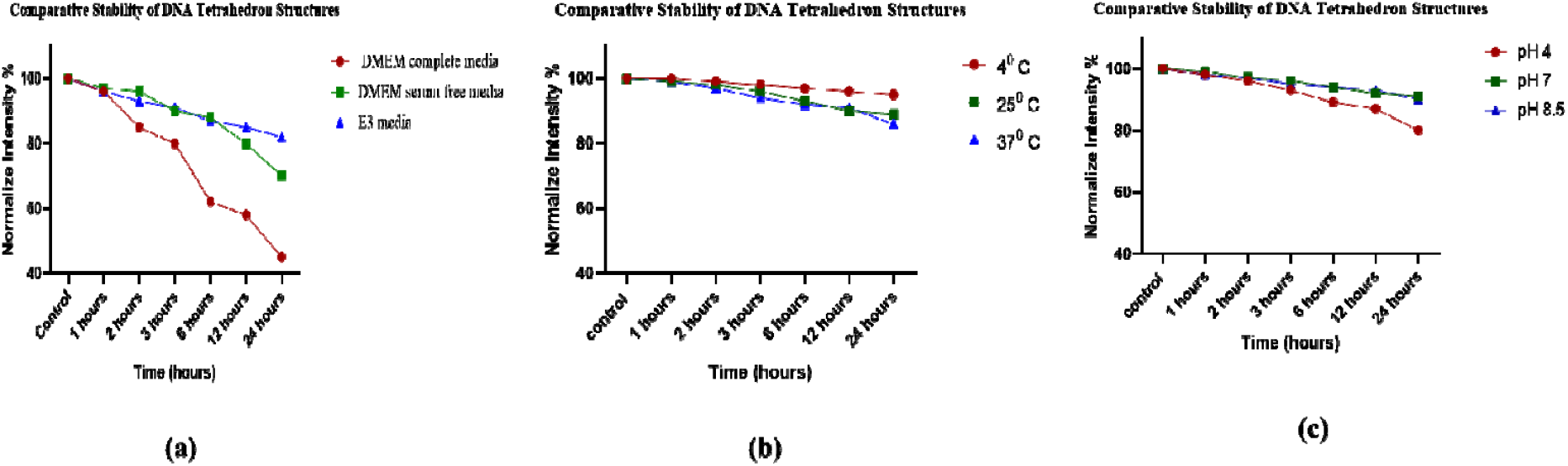
Comparative stability of DNA tetrahedron nanostructures under different conditions over 24 h.(a) Stability in DMEM complete medium, DMEM serum-free medium, and E3 medium.(b) Stability at 4 °C, 25 °C, and 37 °C.(c) Stability at pH 4, pH 7, and pH 8.5.Stability was measured as normalized band intensity (%) at the indicated time points. All samples showed similar initial stability (∼100%). A gradual decrease in stability was observed under all conditions, with the fastest degradation observed in DMEM complete medium, at 37 °C, and at pH 4.

## 4. Discussion

The present study demonstrates that the stability of DNA tetrahedron nanostructures (TDNs) is strongly influenced by biological medium, temperature, and pH conditions. Similar observations have been reported previously, where DNA nanostructures showed reduced stability in serum-containing environments due to nuclease-mediated degradation [16]. In agreement with these reports, our results showed faster degradation in DMEM complete medium, likely because the presence of serum proteins and nucleases promotes the enzymatic breakdown of DNA nanostructures. However, TDNs exhibited comparatively higher stability in serum-free media and E3 medium. E3 medium mainly contains salts and ions required for maintaining osmotic balance, and the absence of serum nucleases may contribute to improved structural stability. Earlier studies have also demonstrated that ionic environments with sufficient salt concentration help preserve DNA nanostructure integrity by stabilizing electrostatic interactions between DNA strands [10] [13]. Our findings therefore support earlier reports describing enhanced stability of DNA nanostructures in ion-balanced and low-nuclease environments. Temperature-dependent analysis further revealed that TDNs remained more stable at 4 °C, whereas increased degradation occurred at 37 °C. Similar temperature-dependent destabilization has been reported for DNA nanostructures, where elevated temperatures enhance molecular motion and enzymatic activity, resulting in accelerated degradation [15]. The comparatively high stability observed at low temperature may be attributed to preservation of Watson–Crick base pairing and reduced nuclease activity. These observations are consistent with previous findings describing improved structural stability of DNA nanostructures under refrigerated conditions.

The pH-dependent stability analysis revealed significant degradation of DNA tetrahedron nanostructures under acidic conditions (pH 4), while comparatively enhanced stability was observed at neutral and mildly alkaline pH conditions (pH 7 and pH 8.5). The reduced stability under acidic conditions may result from proton-induced disruption of hydrogen bonding interactions and destabilization of Watson–Crick base pairing, leading to conformational alterations in the tetrahedral structure. In contrast, neutral and mildly basic pH conditions support duplex stabilization and preservation of structural integrity. Similar pH-responsive behavior in DNA nanostructures has been reported previously in pH-Responsive DNA architectures [44]. Therefore, the present findings agree with earlier studies describing pH-mediated structural destabilization of DNA nanostructures. Overall, the findings of this study are consistent with previously reported studies while also highlighting the comparative stability behavior of DNA tetrahedrons across multiple biologically relevant conditions. Unlike earlier studies primarily focused on structural design or biomedical applications, the present work systematically evaluates the combined influence of media composition, temperature, and pH on TDN stability using normalized PAGE band intensity analysis. These findings suggest that low temperature, neutral to mildly basic pH, and serum-free salt-containing environments such as E3 medium are favorable for maintaining DNA tetrahedron stability and may support their future biomedical and nanotechnological applications [33] [34].

## 5. Conclusion and Future Perspectives

This study provides a systematic evaluation of the stability of DNA tetrahedron nanostructures (TDNs) under biologically relevant conditions, including different media, temperature, and pH. Native PAGE analysis combined with normalized band intensity measurements confirmed successful assembly and enabled comparison of structural stability over 24 h. TDNs showed clear environment-dependent behavior, with serum-containing DMEM complete medium, higher temperature, and acidic pH promoting faster degradation, while serum-free conditions, low temperature, and neutral to mildly basic pH improved structural stability. These findings demonstrate that environmental conditions play a critical role in maintaining TDN integrity and highlight their potential for biomedical and nanotechnology applications.

Future studies should focus on improving the biological stability and functional performance of TDNs through chemical modifications and protective strategies such as PEGylation, phosphorothioate backbone modification, and lipid encapsulation. In addition, advanced characterization approaches including zeta potential analysis, particle size distribution, and structural shape evaluation could provide deeper insight into how physicochemical properties influence nanostructure stability and biological behavior. Further in vivo investigations using suitable model systems, such as zebrafish, may help understand biodistribution, cellular interaction, and long-term stability under physiological conditions. These studies could support the development of optimized DNA nanostructures for applications in targeted drug delivery, biosensing, and responsive therapeutic systems.

## Acknowledgements

The authors express their gratitude to every member of the D.B. study group for their helpful criticism and thorough evaluation of the work. We sincerely thank the Indian Institute of Technology Gandhinagar for its financial and infrastructure support. This work was funded by ANRF-CRG, ANRF-PAIR, GSBTM, CCRH, MoES-STARS.

## Notes

### Competing Interest Statement

The authors have declared no competing interest.

